# Alterations of adult prefrontal circuits induced by early postnatal fluoxetine treatment mediated by 5-HT7 receptors

**DOI:** 10.1101/2023.11.30.569458

**Authors:** Angela Michela De Stasi, Javier Zorrilla de San Martin, Nina Soto, Andrea Aguirre, Jimmy Olusakin, Joana Lourenço, Patricia Gaspar, Alberto Bacci

## Abstract

The prefrontal cortex (PFC) plays a key role in high-level cognitive functions and emotional behaviors, and PFC alterations correlate with different brain disorders including major depression and anxiety. In mice, the first two postnatal weeks represent a critical period of high sensitivity to environmental changes. In this temporal window, serotonin (5- HT) levels regulate the wiring of PFC cortical neurons. Early life insults and postnatal exposure to the selective serotonin reuptake inhibitor fluoxetine (FLX) affect PFC development leading to depressive and anxiety-like phenotypes in adult mice. However, the mechanisms responsible for these dysfunctions remain obscure. We found that postnatal FLX exposure (PNFLX) results in reduced overall firing, and high-frequency bursting of putative pyramidal neurons (PNs) of deep layers of the medial PFC (mPFC) of adult mice in vivo. Ex-vivo, patch-clamp recordings revealed that PNFLX abolished high-frequency firing in a distinct subpopulation of deep-layer mPFC PNs, which transiently express the serotonin transporter SERT. SERT+ and SERT- PNs exhibit distinct morpho-functional properties. Genetic deletion of 5-HT7Rs prevented the PNFLX-induced reduction of PN firing *in vivo* and pharmacological 5-HT7R blockade precluded altered firing of SERT+ PNs *in vitro*. This indicates a pivotal role of this 5-HTR subtype in mediating 5-HT-dependent maturation of PFC circuits that are susceptible to early-life insults. Overall, our results suggest potential novel neurobiological mechanisms, underlying detrimental neurodevelopmental consequences induced by early-life alterations of 5-HT levels.

## Introduction

In the mammalian brain, the prefrontal cortex (PFC) orchestrates several cognitive functions, including working memory, decision-making, cognitive flexibility, as well as social and affective behavior. These executive functions allow understanding and adapting to an ever-changing environment, thus guiding behavior intelligently(1–5). Alterations of prefrontal cortical morphology and functions have been associated with the emergence of mental disorders in humans and in animal models (6,7). In particular, anxiety and depressive disorders affect a significant proportion of the world population (8–10), they represent a major source of disability in humans, and were shown to be associated with alterations of PFC connectivity and function (11–14).

The PFC undergoes a process of refinement during early postnatal development in rodents, when intensive synaptogenesis and spontaneous neuronal firing occur (15). During this critical period of plasticity and vulnerability (16), the serotoninergic system, which includes serotonin (5-HT) receptors and the selective membrane 5-HT transporter SERT, plays a key role in shaping neuronal circuits (15,17–19). Early life insults, such as maternal separation and perinatal administration of selective 5- HT reuptake inhibitors (SSRIs), affect 5-HT levels, leading to negative and permanent effects on the final cortical maturation (20–22). In this context, the risk of developing neuropsychiatric disorders such as anxiety and depression during adulthood is significantly increased (13,23).

In adult mammals, 5-HT is mainly synthetized and released by neurons of the raphe nuclei (24,25) that project to many forebrain regions, including the PFC, and modulate emotional and cognitive behavior (26). 5-HT action is terminated by its reuptake into presynaptic terminals through the selective serotonin transporter SERT, which is mainly expressed in axonal terminals of raphe neurons (27). Importantly, however, it has been shown in rodents that during early developmental stages SERT is transiently expressed in other brain regions, including the neocortex (28,29). In particular, Slc6a4/SERT is transiently expressed by deep-layer pyramidal neurons (PNs) of the medial PFC (mPFC). These SERT+ PNs are mostly cortico-fugal neurons that project outside of the mPFC and target several subcortical areas, including, among others, the mediodorsal and ventromedial nuclei of the thalamus, the dorsal raphe nucleus, ventral tegmental area and the hypothalamus (30).

SERT is the main target of selective serotonin reuptake inhibitors SSRIs (31), which are the most common medications prescribed for both depressive and anxiety disorders (32). SSRIs inhibit 5-HT reuptake, and their chronic administration results in increased 5-HT levels in the brain (33).Fluoxetine (FLX) was the first SSRI approved to be administered in children up to 8 years old to treat depression and compulsive disorder (34,35), and is commonly prescribed to pregnant women suffering perinatal depression (36). However, different studies performed in rodents have demonstrated that while FLX can alleviate depressive symptoms in adults, this drug has paradoxical long-term effects when administered during developmental critical periods (37). Early-life FLX treatment results in anxiety- and depressive-like behaviors, and it impairs fear extinction, learning and memory in adult mice (38–40).

The mechanisms underlying these developmental effects remain poorly understood. Yet, recent transcriptomic, anatomical and behavioral evidence indicates that genetic deletion or pharmacological blockade of 5-HT7Rs are sufficient to rescue the anxiety/depressive phenotype of adult mice, induced by postnatal FLX treatment (PNFLX) (41).

Although PNFLX was shown to produce robust impairments of emotional behaviors in adult animals (37), whether the PNFLX- dependent depressive-like phenotype corresponds to altered activity of the PFC is unknown. In the present study, we used a combination of *in vivo* and *in vitro* approaches to investigate the effects of PNFLX treatment at the network and circuit level in the mPFC of adult mice. We aimed to elucidate the cellular mechanisms leading to the adult emotional behavioral deficits and then evaluate the role of 5-HTR7receptor in the FLX mediated effects.

We found that early postnatal FLX-treatment strongly reduced firing rate and bursting of putative mPFC pyramidal neurons of adult mice, in both anaesthetized and awake conditions. PNFLX specifically affects the firing behavior of SERT+ PNs, which display specific morpho-functional properties as compared with SERT- PNs. We show that in vivo and ex vivo firing alterations of PNs induced by PNFLX were prevented by genetic deletion or pharmacological blockade of 5-HT7Rs, identifying a new potential target to treat 5-HT-dependent emotional behavioral dysfunctions induced by early-life stress.

## Results

### PNFLX results in decreased firing of deep-layer PFC putative pyramidal neurons in anaesthetized adult mice

To investigate the PFC network consequences of early-life FLX administration, we performed *in vivo* electrophysiological recordings in the PFC of adult mice that were subject to PNFLX (Fig. 1a). Mouse pups where fed with either control sucrose solution or FLX (10% in 2% sucrose solution; See Methods) daily from P2 to P14. This protocol was shown to induce robust alterations of emotional behaviors in adult mice (>P90) (38,41). Therefore, using cell-attached (juxtacellular) recordings, we measured single-unit activity from deep-layers (L5b-L6) of the prelimbic and infralimbic region of the PFC in head-fixed, anesthetized mice. A second glass electrode was used to record the nearby local field potential (LFP) (Fig. 1 A, B). Juxtacellular blind recordings were done from putative pyramidal neurons (PNs), which represent the majority of neurons in the mouse PFC (42), although we cannot exclude that, our datasets include some inhibitory interneurons in both conditions. Putative PN firing in anesthetized mice exhibited bouts of spiking activity mainly occurring during UP states of the LFP (Fig. 1B). In addition, high-frequency bursts were commonly observed (19/23 and 23/27 PNs in CTR and PNFLX, respectively) and characterized by a prominent peak in the inter-spike interval (ISI) distributions, corresponding to the highest instantaneous firing frequencies (Fig. 1c; see Methods). We found that PNFLX produced a ∼50% reduction of overall putative PN firing rate (5.52 ± 1.40 vs. 2.76 ± 0.93 Hz, CTR vs. PNFLX, n = 23 and 27, respectively; p < 0.05, Mann-Whitney U-test; Fig. 1 b, d). Interestingly, we found that PNFLX affected the ability of putative PNs to fire bursts. Indeed, in control condition, the distribution curves of inter-spike intervals (ISIs) exhibited a prominent peak at short ISIs, corresponding to high-frequency firing. This peak was much smaller in recordings from PNFLX-treated mice (Fig. 1c). In general, PNFLX treatment reduced burst occurrence (burst rate: 0.3 ± 0.06 vs. 0.13 ± 0.05 Hz; CTR vs. PNFLX, n = 19 and 23, respectively; p < 0.01, Mann-Whitney U-test; Fig. 1 b, d). Bursts lasted significantly longer in PNFLX mice (0.12 ± 0.06 vs. 0.32 ± 0.1 s in CTR vs. PNFLX; n = 19 and 22 neurons and N = 8 and 9 mice, respectively; p < 0.05, Mann-Whitney U-test; Fig. 1 b, d), with a similar number of spikes per burst (Fig. 1c; p > 0.05; n = 19 and 22, respectively; Mann-Whitney U-test).

**FIGURE 1.**
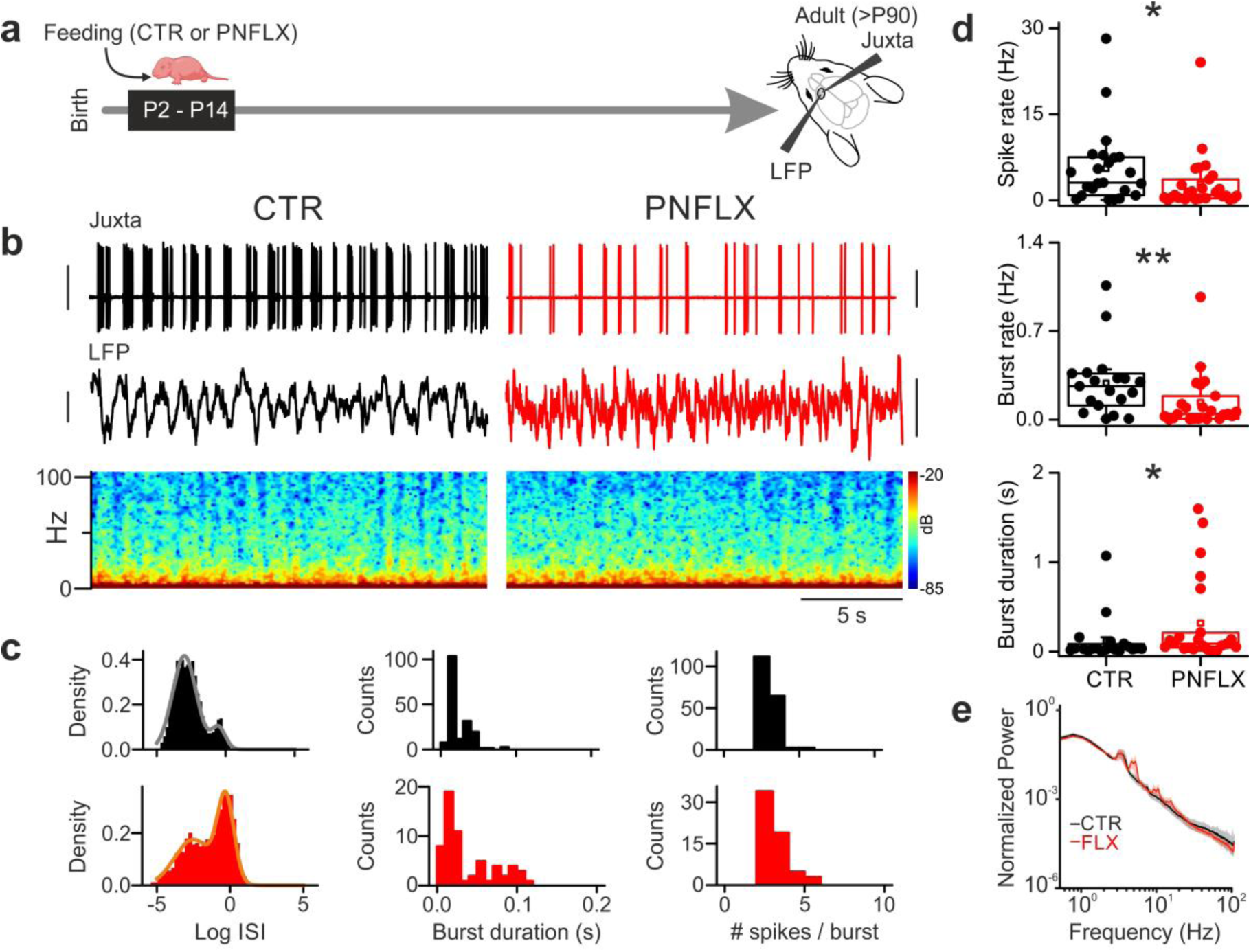
Cortical network dynamics in anaesthetized CTR and FLX treated mice. **a:** Schematic representation of the perinatal fluoxetine protocol (PNFLX) and experimental configuration. Pups were fed from P2 to P14 with either fluoxetine (PFLX group) in 2% sucrose solution, or 2% sucrose solution alone (CTR group). Electrophysiological recordings of spontaneous firing (juxtacellular) and LFP activity were performed in the PFC of adult mice. **b:** Representative traces of in vivo extracellular recordings in urethane-anaesthetized CTR (left black) and PFLX (right-red) mice. Top: loose-patch, single unit spikes activity. Bottom: Local field potential (LFP) population activity and its relative spectrogram. Vertical scale bars: 5 mV and 0.5 mV for juxtacellular and LFP recordings, respectively. **c:** Histograms illustrating the inter-spike interval distribution (left), burst duration (center) and number of spikes within burst (right) of the cell shown in B. **d:** Population plots of spike rate (top), burst rate (middle) and burst duration (bottom) in CTR (black dots) and PFLX (red dots) animals. **e:** Average normalized LFP power spectral densities for PFLX and CTR mice

We found no alterations of overall LFP activity in all frequency bands, as indicated by overlapping average power-spectrum density curves in both control and PNFLX-treated animals (Fig. 1b, e).

Overall, our results indicate that early-life FLX treatment induces a strong reduction of firing frequency in deep-layer putative PNs of the adult mouse PFC with no apparent alterations of basic network activity. Reduced PN firing is likely due to a PNFLX-mediated reduction of high-frequency burst firing.

### PNFLX results in decreased firing of deep-layer PFC putative pyramidal neurons in awake adult mice

Deep anesthesia, including that induced by urethane, strongly affects cortical dynamics by inducing various forms of highly synchronous activity (43–45). It is therefore possible that the PNFLX-induced effects described in Fig. 1 are amplified or present only in anesthetized animals. To rule out this possibility, we performed the same experiments but in awake, head-fixed mice (Fig. 2a; see Methods). We found that spiking activity was strongly reduced in PNFLX mice, as compared to their control littermates (Fig. 2b, d; firing rate: 2.7 ± 0.5 vs. 0.4 ± 0.1 Hz, CTR vs. PNFLX n = 20 and 13 neurons and N = 4 and 3 mice, respectively; p < 0.001, Mann-Whitney U-test). Likewise, bursting activity was strongly diminished in PNFLX mice (Fig. 2b-d; burst rate: 0.1 ± 0.03 vs. 0.02 ± 0.01 Hz, CTR vs. PNFLX n = 20 and 13 neurons and N = 4 and 3 mice, respectively; p < 0.001, Mann-Whitney U-test). Despite occurring at lower frequencies, burst duration was not different between control and PNFLX mice (Fig. 2c, d; burst duration: 0.07 ± 0.02 vs. 0.2 ± 0.1 s, CTR vs. PNFLX n = 20 and 13 neurons and N = 4 and 3 mice, respectively; p > 0.05, Mann-Whitney U-test). In awake mice, we observed a more desynchronized LFP, lacking prominent UP and DOWN states typical of alert states. Similarly to the anesthetized condition, LFP was unaffected by PNFLX treatment (Fig. 2b, e).

**FIGURE 2.**
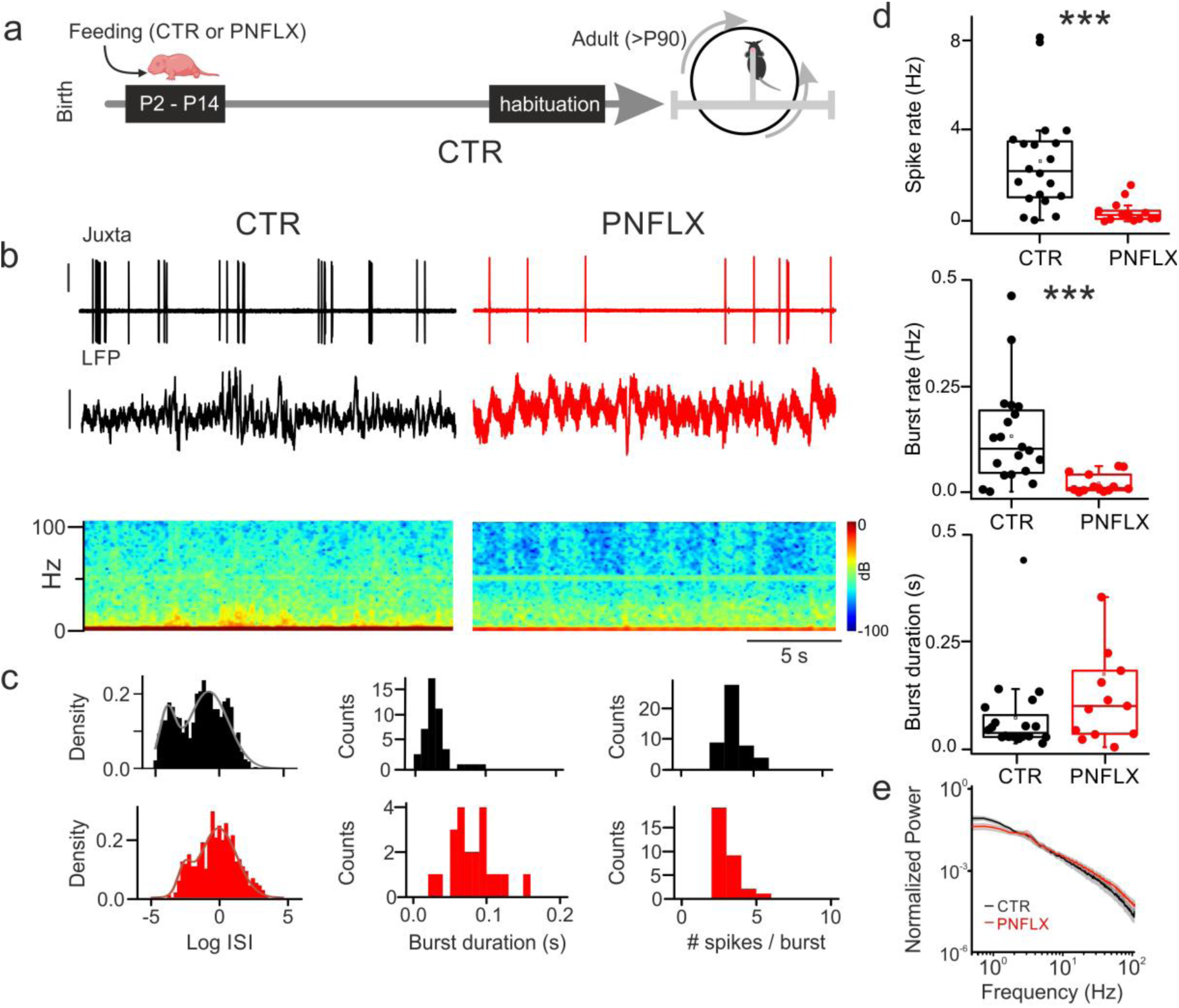
Cortical network dynamics in awake head-fix CTR and FLX treated mice. a: Schematic representation of the perinatal fluoxetine protocol (PNFLX) and experimental configuration. Electrophysiological recordings were then performed in awake head restrained adult mice that were allowed to move in a floating cage apparatus. b: Representative traces of in vivo extracellular recordings in urethane-anaesthetized CTR (left black) and PFLX (right-red) mice. Top: loose-patch, single unit spikes activity. Bottom: Local field potential (LFP) population activity and its relative spectrogram. c: Histograms illustrating the inter-spike interval distribution (left), burst duration (center) and number of spikes within burst (right) of the cell shown in B. d: Population plots of spike rate (top), burst rate (middle) and burst duration (bottom) in CTR (black dots) and PFLX (red dots) animals. e: Average normalized LFP power spectral densities for PFLX and CTR miceCTR mice

These results indicate that PNFLX causes a strong reduction of putative PN firing and bursting frequencies in adult mice that are not mediated by anesthesia.

### PNFLX-mediated reduction of *in vivo* firing requires 5-HT7 receptors

5-HT7Rs were shown to be highly expressed during development where they were found to play a role in axon growth and synaptogenesis (46–48). Moreover, this specific 5-HTR subunit is transiently co-expressed with SERT by PFC neurons of the PFC. Accordingly, genetic deletion of 5-HT7Rs prevents the detrimental emotional behavioral effects caused by FLX exposure during early postnatal life, indicating a new role for 5-HTR7 in the postnatal maturation of prefrontal circuits (41). It is therefore possible that the PNFLX-mediated reduction of PN firing acts through the same 5-HTR subunit. We therefore recorded from putative PNs in 5-HT7 KO mice and their WT littermates, in anesthetized control (sucrose-fed) and PNFLX-treated animals. We found that PNFLX induced a significant, ∼50% reduction of firing rates in WT animals, also in this mouse line (firing rate: 2.57 ± 0.42 and 1.23 ± 0.21 Hz in WT-CTR and WT-PNFLX, respectively; n = 28 cells from 5 mice and n = 36 cells from 5 mice, respectively; p < 0.05, Kruskal–Wallis one-way ANOVA, followed by Dunn’s multiple comparisons; Fig. 3ac).

**FIGURE 3.**
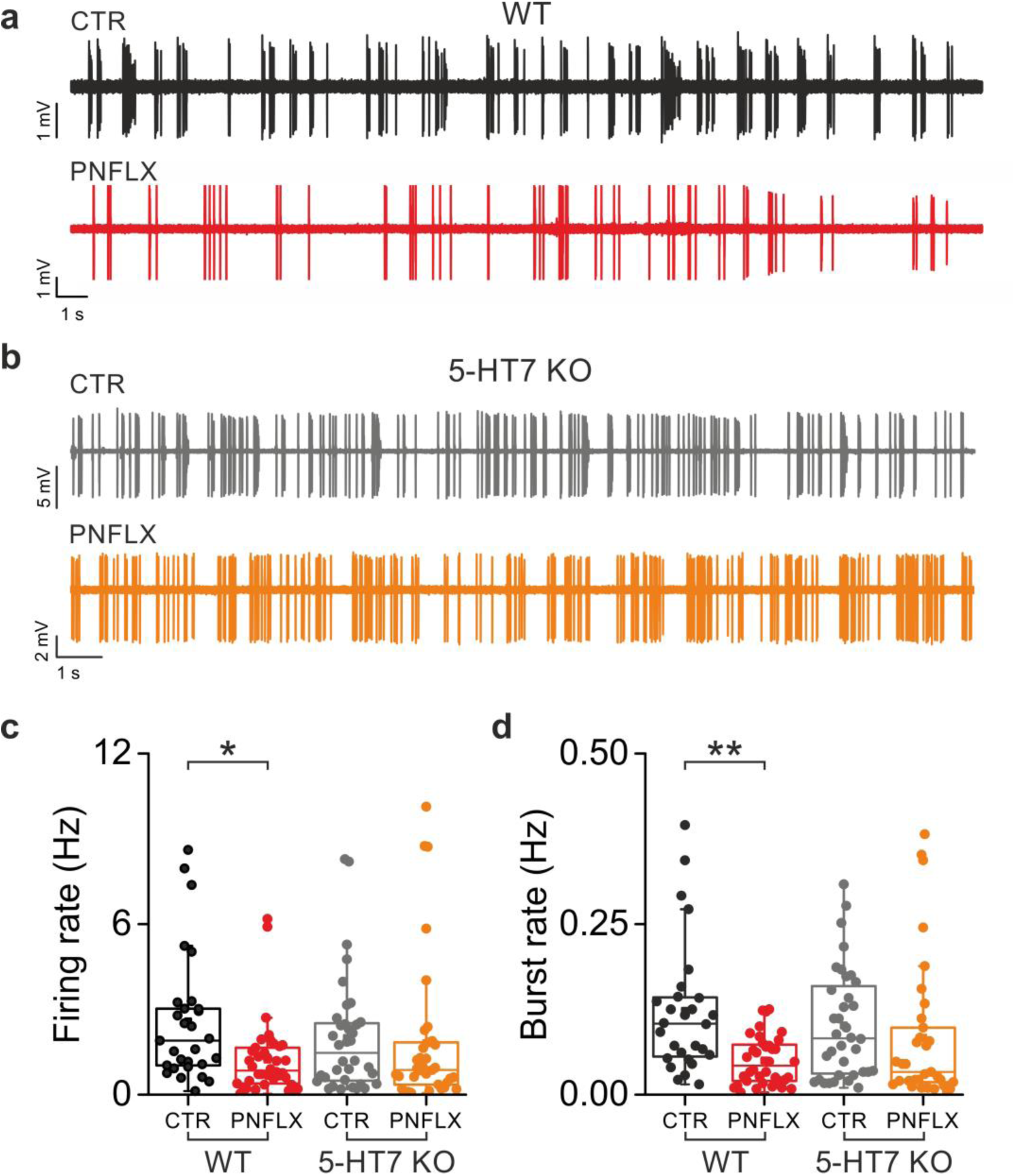
PNFLX does not affect in vivo firing of PFC putative pyramidal neurons in 5-HT7 KO mice. **a:** Representative traces of loose-patch, single unit recordings from two putative PFC pyramidal neurons recorded in a 5-HT7 WT CTR mouse (black trace, top) and in a 5-HT7 WT PFLX (red trace, bottom) mouse. **b:** As in A, but in a 5-HT7 KO mouse (gray trace: CTR; orange trace: PNFLX). **c:** Population plots showing the firing and burst rates in 5-HT7 WT CTR (black dots), 5-HT7 WT PFLX (red dots), 5-HT7 KO CTR (grey dots) and 5-HT7 KO PFLX (orange dots) group.

Conversely, firing activity was unaffected by PNFLX treatment in 5-HT7 KO mice (firing rate 1.99 ± 0.34 and 1.81 ± 0.43 Hz, 5HT7-KO-CTR and 5HT7-KO PNFLX mice, respectively; n=36 cells from 7 mice and 33 cells from 8 mice, respectively; p > 0.05, Kruskal–Wallis one-way ANOVA, followed by Dunn’s multiple comparisons; Fig. 3ac). Likewise, high-frequency burst rate was strongly affected by PNFLX in WT, but not 5-HT7 KO animals (p < 0.01 and p > 0.05 in WT and 5-HT7 KO, respectively, Kruskal–Wallis one-way ANOVA, followed by Dunn’s multiple comparisons; Table 1; Fig. 3b, d).

**Table 1.**

|  | WT |  | 5-HT7 KO |  |
| --- | --- | --- | --- | --- |
|  | CTR<br>(n = 28; N = 5) | PNFLX<br>(n = 36; N = 5) | CTR<br>(n = 36; N = 7) | PNFLX<br>(n = 33; N = 8) |
| Firing rate (Hz) | 2.6 ± 0.4 | 1.2 ± 0.2* | 2.0 ± 0.3 | 1.8 ± 0.4 |
| Burst rate (Hz) | 0.12 ± 0.02 | 0.05 ± 0.01** | 0.1 ± 0.02 | 0.08 ± 0.02 |
| Burst duration (s) | 0.05 ± 0.01 | 0.1 ± 0.03 | 0.1 ± 0.02 | 0.2 ± 0.03 |

Mean ± SEM; \*P < 0.05; \*\*P < 0.01, Mann-Whitney U-test.

These results indicate that increased 5-HT levels induced by early-life SSRI exposure exert detrimental effects in neuronal firing dynamics of the PFC in adult mice, by acting on 5-HT7Rs.

### PNFLX treatment affects the firing dynamics of a specific subtype of deep-layer SERT+ PNs via 5- HT7Rs

In P0-P10 mice, the 5-HT transporter SERT is transiently expressed in a subset of layer 5-6 pyramidal neurons of PFC. Importantly the PFC neurons that transiently express SERT were found to play a crucial role in the depression and anxiety-like symptoms that result from PNFLX exposure. SERT+ neurons are dense in deep layers of the PFC, and they identify a molecularly defined subset of PNs (30). Therefore, it is possible that reduced firing of PNFLX-treated mice observed *in vivo* was restricted to SERT+ PNs. To test this hypothesis, we crossed SERT-Cre mice with a tdTomato reporter line to label SERT+ PNs (Fig. 4a, b; see Methods). P2-14 Sert-Cre::TdTomato mice were fed with either sucrose (CTR) or sucrose and fluoxetine (PNFLX; Fig. 4c). We then obtained acute brain slices from adult mice, and performed whole-cell, current-clamp recordings from either SERT+ or SERT- PNs (Fig. 4b, c). We constructed input-out curves by injecting 800 ms-long DC currents and recorded the corresponding firing behavior. In control conditions, we found that SERT+ PNs were characterized by higher firing frequency than SERT- PNs of layers 5-6 (Fig. 4d, e; maximum firing frequency: 69.72 ± 5.70 vs. 34.38 ± 6.84 Hz SERT+ vs. SERT- PNs; p < 0.001 2way ANOVA, followed by Tuckey’s multiple comparisons, n = 9 SERT+ CTR; n=8 SERT- CTR). In addition, SERT+ PNs exhibited a significantly less adapting firing behavior than SERT- PNs (Suppl Fig. 1). Interestingly, PNFLX treatment resulted in disruption of high-frequency firing, selectively in SERT+ PNs, as firing dynamics of SERT- PNs were unaffected by PNFLX (Fig. 4d, e; maximum firing frequency: 69.7 ± 5.70 vs. 33.25 ± 6.05 Hz in SERT+ PNs, CTR vs. PNFLX, respectively; p < 0.01, 2way ANOVA, followed by Tuckey’s multiple comparisons, n = 9 and 15 SERT+ CTR and PNFLX, respectively; 34.38 ± 6.84 vs. 32.78 ± 3.27 Hz, SERT- PNs, CTR vs. PNFLX, respectively; p > 0.05, 2way ANOVA, followed by Tuckey’s multiple comparisons, n = 8 and 9 SERT- CTR and PNFLX, respectively). We analyzed passive properties of SERT+ and SERT- PNs in both conditions and did not find any significant differences in membrane resistance, resting membrane potential, rheobase and sag (Suppl. Fig. 1). Similarly, we did not find any differences in action-potential threshold, half-width, amplitude, max depolarization and hyperpolarization speed in both SERT+ and SERT- PNs in control and PNFLX (Suppl. Fig. 2).

**FIGURE 4:**
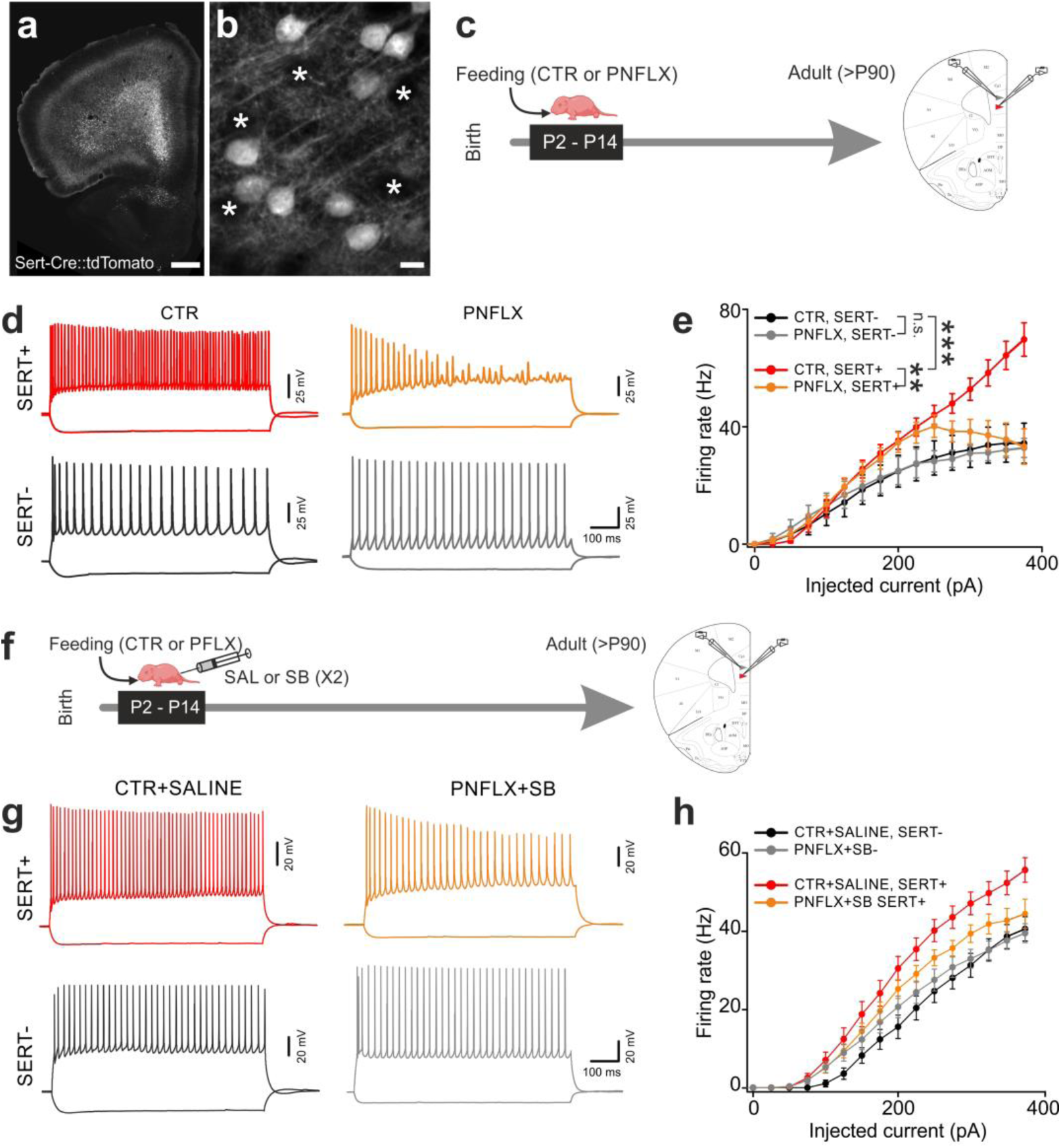
SERT Positive specific dysfunction induced by PNFLX protocol is rescue by administration of 5-HT7 antagonist in PFC brain slices. a: Confocal fluorescence micrograph illustrating a section of the PFC of a SERT-Cre::tdTomato adult mouse. Note the dense labeling of deep-layer tdTomato-positive neurons. Scale bar: 500 µm b: Close-up of the micrograph of A showing deep-layer, fluorescent SERT+ cells and shadows of SERT- cells (asterisks). Scale bar: 10 µm. c: Scheme of the experimental procedure. PNFLX protocol was followed by patch-clamp recordings in acute brain slices obtained from adult mice. d: Representative current-clamp traces showing the response to 0.8 s-long current injections from SERT+ and SERT- PNs. Red and orange traces: SERT+ PNs in CTR and PNFLX, respectively. Black and gray traces: SERT- PNs in CTR and PNFLX, respectively. current injections: from -100 to +375 pA in all cases. e: Firing rate vs. current injections in all cases. Same colors as in D. *P < 0.05; **P < 0.01; ***P < 0.001. f: Scheme of the experimental procedure. Same as in C, except that during PNFLX protocol mice received concomitant intraperitoneal injections of saline solution or saline solution containing the 5-HT7 antagonist (SB269970) twice a day. g-h: Same as in d-e but in the presence of SB269970 (SB).

To test whether 5-HT7Rs are involved in disruption of SERT+ PN high-frequency firing, we repeated the same experiments in adult mice that were fed with either sucrose or FLX at P2-P14 and received pharmacological treatment with the selective 5-HT7R antagonist, (2R)-1-[(3-hydroxyphenyl)sulfonyl]- 2-[2-(4-methyl-1-piperidinyl)ethyl]pyrrolidine (SB 269970 hydrochloride, here referred to as SB; Fig. 4F) (41,49). To make sure that 5-HT7Rs were blocked during the entire duration of PNFLX protocol, mouse pups received two SB or vehicle (saline) injections per day (see Methods). We repeated the same current clamp experiments in acute brain slices as described above. We found that pharmacological blockade of 5-HT7Rs during PFLX treatment prevented the disruption of high-frequency firing of SERT+ PNs (Fig. 4g, h; n = 11 and 7, SERT+ and SERT- neurons, respectively, from N =4, CTR SALINE mice respectively; n = 35 and 18 SERT+ and SERT- neurons, respectively, from N = 8 PNFLX SB mice). No effect of SB was detected on the firing dynamics of SERT- PNs.

Altogether, these experiments reveal that SERT+ and SERT- PNs belong to two different sub-populations of deep-layer principal neurons that can be identified by their biophysical properties. Importantly, PNFLX treatment selectively prevented SERT+ PNs from firing at high frequencies, without affecting the intrinsic excitability of SERT- PNs.

### PNFLX does not affect the morphology of deep-layer, SERT+ PNs of the PFC

We found that SERT+ PNs exhibit a specific firing behavior characterized by the ability of firing at relatively high frequency (Fig. 4). Deep-layer PNs of the PFC were described as belonging to at least two different subclasses (50,51). Moreover, there is some evidence that interfering with the 5-HT system affects the arborization of prelimbic and infralimbic PNs (37). We wanted to test if SERT+ and SERT- PNs could also identify two morphologically distinct principal neuron subclass, and whether PNFLX treatment alters their dendritic morphology. To this end, we fed SERT-Cre::tdTomato mice with either sucrose (CTR) or sucrose and FLX (PNFLX) at P2-P14. We then obtained acute PFC slices from adult mice and performed long lasting (>30 min) whole-cell recordings from both SERT+ and SERT- PNs, including in the intracellular solution high concentration (5 mg/mL) of biocytin (Fig. 5a, b). After histological processing, biocytin-positive SERT+ and SERT- neurons were reconstructed and the complexity of their basal and apical dendrites was quantified, using Scholl analysis (Fig. 5c-d). Basal dendrites did not exhibit any significant difference across cell types and treatments (Fig. 5c-d). Interestingly, however, the apical dendrites were significantly more branched in SERT+ than SERT- PNs (Fig. 5c-d; p < 0.005; Mixed-effects model (restricted maximum likelihood, REML), followed by Tuckey’s multiple comparisons; n = 8 and 6, SERT+, CTR and SERT-, CTR, respectively). PNFLX treatment did not significantly alter the overall dendritic arborization of SERT+ PNs (Fig. 5c-d; p > 0.05; Mixed-effects model (restricted maximum likelihood, REML), followed by Tuckey’s multiple comparisons; n = 8 and 8, SERT+, CTR and SERT+, PNFLX, respectively). However, PNFLX treatment had a small albeit significant effect on SERT- PNs (Fig. 5c-d; p < 0.05; Mixed-effects model (restricted maximum likelihood, REML), followed by Tuckey’s multiple comparisons; n = 6 and 8, SERT-, CTR and SERT-, PNFLX, respectively).

**FIGURE 5.**
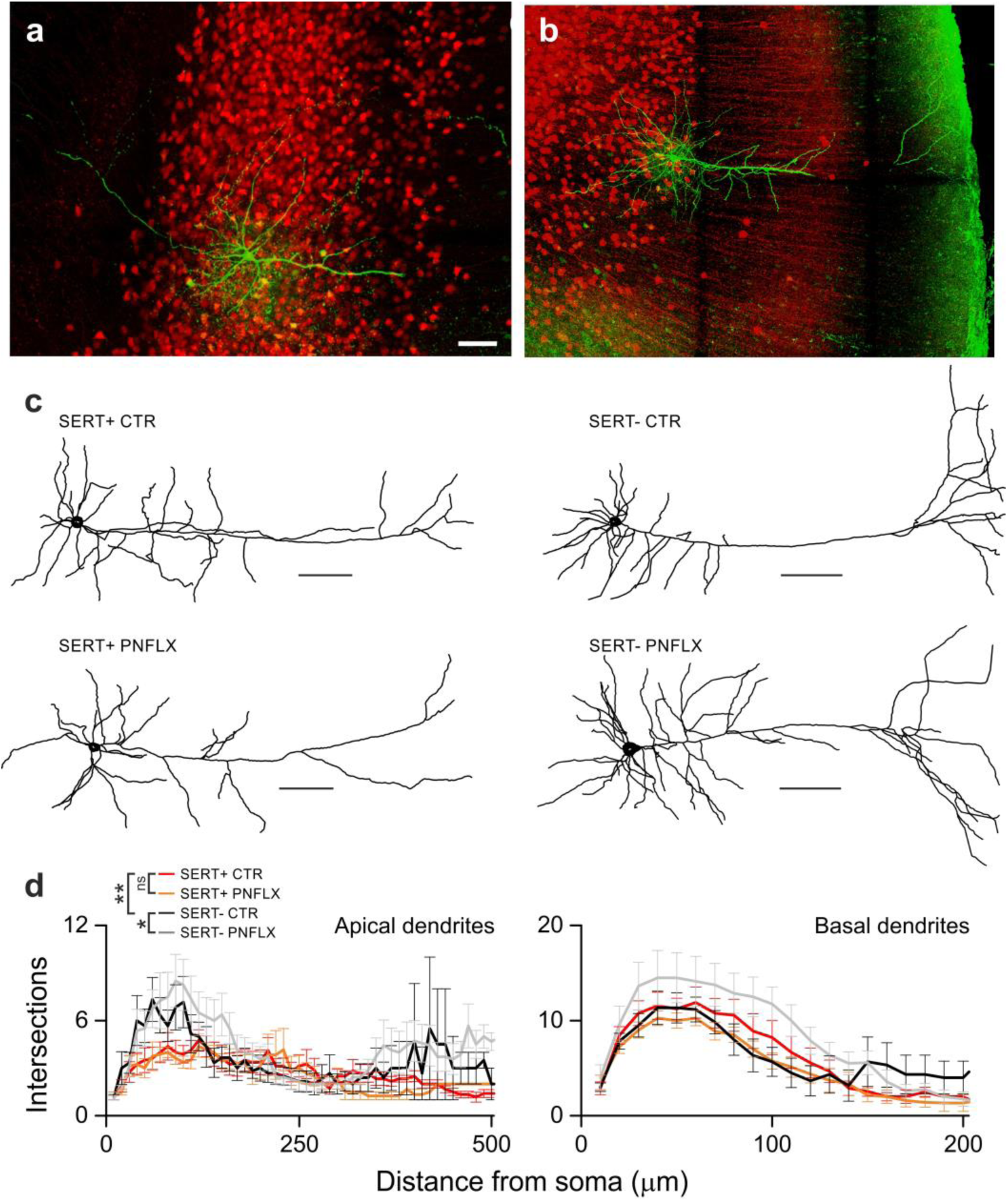
Morphological characterization of SERT Positive and SERT Negative CTR and PFLX pyramidal neurons in deep layers of adult PFC. a-b: Fluorescent micrographs showing a SERT+ (left) and a SERT- (right) PN filled with biocytin and counterstained with green, fluorescent streptavidin. Neurons were filled in a SERT-Cre::TdTomato mouse. Red fluorescence indicates SERT+ PNs. Scale bar 50 µm. c: Representative somatodendritic reconstructions of biocytin-filled SERT+ and SERT- PNs in CTR and PNFLX. Scale bar 100 µm. d: Number of intersections plotted as a function of the distance from soma for basal (lef) and apical (right) dendrites for the two cell types (SERT+ and SERT-), for both treatments (CTR and PNFLX). The yellow box corresponds to the data points that are significantly different between SERT+ and SERT- PNs (* : P<0.005).

Overall, SERT+ PNs were morphologically less complex than SERT- PNs, whose apical dendrites tufted extensively in layer 1. These experiments indicate that, in addition to their specific excitability, SERT+ PNs also exhibit peculiar morphological features. Importantly, PNFLX treatment does not overall alter adult deep-layer PN morphology.

### PNFLX does not affect global spontaneous excitatory and inhibitory synaptic transmission in adult deep-layer SERT+ and SERT- PNs

Excitation-to-inhibition balance onto principal cortical PNs has been ascribed as a prominent pathophysiological mechanism underlying the emergence of several psychiatric diseases, including depression and anxiety (52,53). Indeed, the marked reduction of PN firing induced by PNFLX treatment and observed *in vivo* (Figs. 1-3) suggests that adult PFC cortical circuits might suffer from alterations of either glutamatergic and/or GABAergic neurotransmission. To test this, we performed whole-cell, voltage-clamp recordings from SERT+ and SERT- PNs in acute PFC slices obtained from adult SERT- Cre:tdTomato mice that were fed with control solution (sucrose) or FLX at P2-P14. We recorded spontaneous glutamatergic and GABAergic neurotransmission as a proxy for global synaptic excitation and inhibition and thus probe general pre- and/or postsynaptic alterations induced by PNFLX treatment in SERT+ and SERT- PNs. We isolated glutamatergic spontaneous excitatory postsynaptic currents (specs) by using a low-chloride intracellular solution (see Methods) and by blocking GABAergic neurotransmission using the GABA_A_R antagonist gabazine (10 µM; Fig. 6A). We found that sEPSC frequency and amplitude were similar across cell types and treatment (Fig. 6a, b; n=12 and 12 SERT+ and SERT- cells respectively in N= 8 CTR mice; n=10 and 11 SERT+ and SERT- cells respectively in N= 6 PNFLX mice). Spontaneous, GABAergic inhibitory postsynaptic currents (sIPSCs) were isolated by using a high-chloride-containing intracellular solution and in the continuous presence of the ionotropic glutamate receptor antagonist DNQX (10 µM) and AP5 (100 µM) (Fig. 5C). Similarly to sEPSCs, PNFLX treatment did not affect sIPSC frequency and amplitude in both SERT+ and SERT- PNs (Fig. 5c, d; n= 8 and 10 SERT+ and SERT- cells respectively in N= 4 CTR mice; n=16 and 15 SERT+ and SERT- cells respectively, in N= 5 PNFLX mice).

**FIGURE 6.**
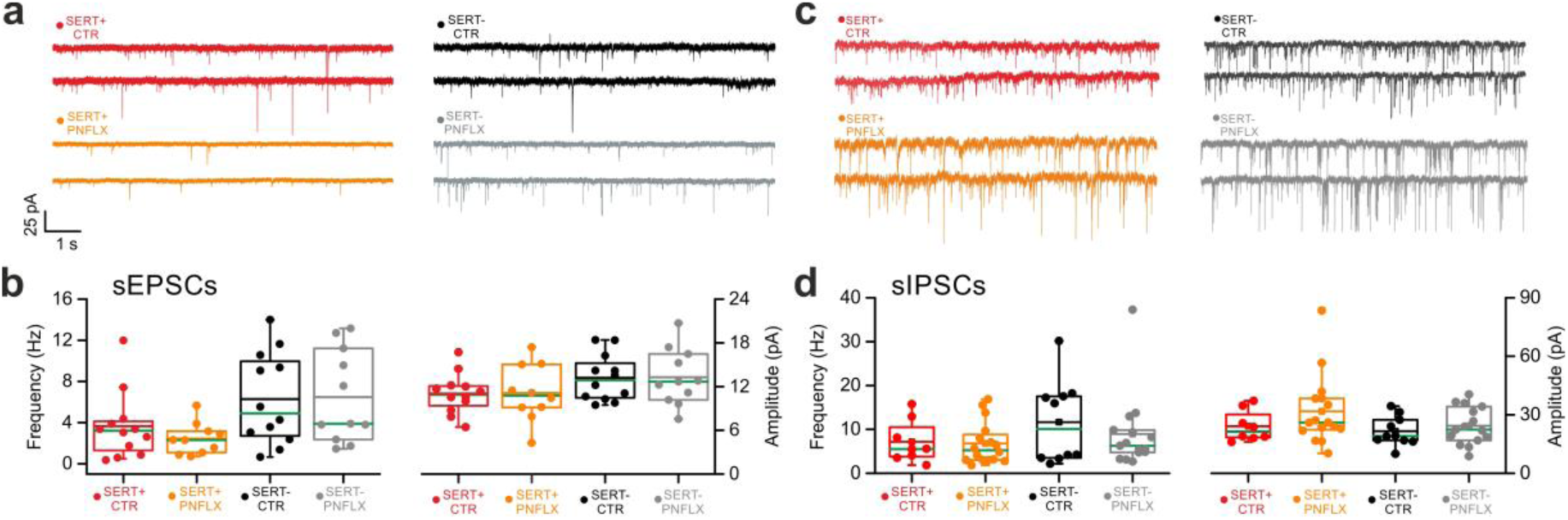
Spontaneous excitatory (EPSCs) and inhibitory (IPSCs) activities recorded in SERT+ and SERT- PNs in CTR and PFLX adult mice. **a:** Representative voltage-clamp traces of glutamatergic sEPSCs recorded from PFC PNs of adult CTR and PNFLX-treated mice. **b:** Population data of sEPSC frequency (left) and amplitude (right) in SERT+ and SERT- PNs in CTR and PNFLX-treated mice. Green and black lines indicate median and mean values. resepctively. **c-d:** Same as in a-b but for GABAergic sIPSCS, isolated using a high-chloride intracellular solution and in the continuous presence of ionotropic glutamate receptor antagonists in the extracellular superfusate

Overall, we did not detect major differences of global excitatory and inhibitory neurotransmission between SERT+ and SERT- PNs. Moreover, we did not detect any alteration of both spontaneous glutamatergic and GABAergic neurotransmission induced by PNFLX treatment in both cell types. These experiments indicate that PNFLX does not alter the basic properties of excitatory and inhibitory neurotransmission.

## Discussion

In this study, we used PNFLX treatment to mimic early-life-induced anxiety and depression in adults. In rodents, PNFLX protocol yields such a robust anxious and depressive phenotype in adults that it has become a well-established model used to study the pathophysiological mechanisms of depression with a developmental component (37). In addition to being an early-life insult, PNFLX treatment has also an important translational aspect as SSRIs are commonly prescribed to pregnant and lactating women as well to young children (54–56). Robust mouse models of psychiatric diseases give access to a powerful genetic toolbox to study cortical network dysfunctions and alterations of distinct elements of the cortical circuit with unprecedented detail. We found that early-life FLX administration during a specific 5HT sensitive period strongly decreases putative pyramidal neurons spiking activity without affecting the global network dynamics through 5-HT7Rs. We found that a specific subclass of deep-layer PNs exhibits significant alterations of intrinsic excitability. These neurons can be distinguished morphologically and electrophysiologically and transiently express SERT. PNFLX dependent alterations were caused by an overactivation of 5-HT7R, as they were prevented when FLX was administered concomitantly with a 5-HT7R antagonist and did not occur in 5-HT7-KO mice.

We found that deep-layer putative PNs exhibit high-frequency burst firing. Bursts, defined as bouts of high-frequency activity (see Methods), occurred less frequently in PNFLX-treated mice, likely contributing to the overall reduction of firing frequency. Interestingly, SERT+ PNs could not sustain firing activity at frequencies higher than ∼40 Hz. This suggests that a large majority of neurons sampled in our in vivo recordings were SERT+. This is consistent with the high density of SERT+ PNs in deep layers of the PFC, especially L6 (see Figs. 4 and 5) (41). In addition, the temporal window of SERT postnatal expression by these PNs (P0-P10) is consistent with the PNFLX pharmacological treatment. It is important to note, however, that the alterations of SERT+ PN firing that we observed in our slice recordings is likely not the only cause of the strong reduction of spiking activity that we measured in our *in vivo* recordings. Several other factors might account for this, including changes in feed-forward inhibition triggered by long-range afferents such as, for example, those originating in the medial thalamus, which play a major role in shaping the activity of PFC circuits involved in cognitive and emotional behaviors (57–59)

Although reducing PN firing rates, PNFLX administration during a specific 5HT sensitive period does not affect the global network dynamics, as measured in our LFP recordings. This could be due to the fact the LFP signal spreads across different layers and originates from a wide cortical area. The lack of differences in the LFP power spectrum in control vs. PNFLX-treated mice, especially in the higher-frequency bands, suggests that inhibitory circuits might be overall unaffected in this model. Indeed, we found no global changes in spontaneous inhibitory neurotransmission. This contrasts with other studies indicating that disruptions of emotional behavior in a genetic animal model of schizophrenia correlates with marked alterations of theta and gamma oscillations, paralleled by dysfunctions of PV-cell-mediated synaptic properties (60). This apparent discrepancy indicates that comparable dysfunctions of emotional behavior can emerge from either major inhibitory circuit alterations (induced by a genetic cause that is persistent throughout life) or specific neurotransmitter imbalances occurring during a specific sensitive developmental window. We cannot rule out the possibility that intrinsic firing properties of specific interneurons subclasses and action potential-evoked inhibitory neurotransmission is affected by PNFLX treatment.

Since general anesthesia is known to affect brain rhythms and cellular activities (43,45) we repeated the same experiments in awake, head-restrained mice that were free to locomote in a floating cage. Even in these conditions, we detected a major reduction in firing frequency and burst rate in PNFLX-treated mice with no overall change in the power spectrum of the LFP, indicating that PNFLX treatment induces a strong PFC network phenotype in the adult that is independent from a specific brain state.

Previous evidence indicates that 5HT-7R signaling modulates neuronal motility and morphology, dendritic spines, and synaptogenesis in different brain areas, suggesting a key role for this receptor in mediating a 5-HT trophic-like mechanism during development(61,62). Moreover, mice lacking 5-HT7Rs did not display deficits in emotional behaviors induced by PNFLX treatment. Accordingly, we found that PNFLX-treatment of 5-HT7 KO mice was not accompanied by a significant reduction of firing frequency and burst rates of deep-layer PNs of the PFC. This well agrees with high mRNA expression of 5-HT7Rs in deep layers of the PFC in the first postnatal week, and co-localization with SERT+ PNs(41).The significant reduction detected in WT littermates indicates that the susceptibility to PNFLX-treatment persists also in this different genetic mouse background. Overall, our findings suggest that increased levels of serotonin induced by PNFLX treatment may over-activate 5HTR-7Rs, leading to hypoactivity of pyramidal neurons.

To date, at least 15 5HTRs have been described (18). All of them display specificity of temporal and spatial expression patterns (37), and 5-HT7 is among the earliest to be expressed by deep-layer neurons of the PFC (63). Although we cannot exclude additional and/or synergistic roles of other 5- HTR subtypes, our experiments indicate a central role of 5-HT7Rs in altered network activity. Thus, the 5-HT7-dependent alterations of mPFC PN firing that we describe here can, at least in part, explain the 5-HT7-dependent emotional behavioral deficits induced by PNFLX treatment (37,41). Likewise, despite SB269970 was described to be selective for 5-HT7Rs (64,65), we cannot rule out the possibility of undocumented off-target effects of this compound. However behavioral (30,41) and electrophysiological experiments (this study) suggest a convergence with genetic removal of 5-HT7Rs, underlying the central role played by this receptor subclass.

The reduced firing frequency and burst occurrence in vivo, detected in PNFLX-treated mice, correlated with alterations of firing dynamics recorded in acute brain slices, selectively in SERT+ PNs. As previously shown, these neurons are dense in the deep layers of the PFC, especially in L6 (30). Here we describe that SERT+ PNs represent a specific PN subtype as compared to intermingled SERT- neurons. SERT+ PNs fire at much higher frequency with lower adaptation, and they exhibit less branched apical dendritic trees. The inability to sustain high-frequency firing of SERT+ PNs of PNFLX treated mice could be in principle due to alterations of intrinsic membrane properties and/or differences in the expression of specific ion channels shaping the waveform of action potentials. We did not find changes in passive properties nor in action potential waveform. Lack of changes in membrane resistance and rheobase is consistent with the unaltered slope of the initial part of the f-i curve (Fig. 4). Breakdown of firing at frequencies higher than 40 Hz could be due to altered expression of specific Ca^2+^-dependent K^+^ channels that regulate the ability of firing repeatedly in neurons (66,67). Future studies will be necessary to identify the specific ionic conductance(s) responsible for high-frequency firing in SERT+ PNs that is modified by early-life treatment with FLX. Interestingly, this high-frequency firing breakdown in SERT+ PNs was recovered in mice, whose PNFLX-treatment was accompanied by administration of SB269970, a selective 5-HT7R antagonist (68). Pharmacological 5- HT7R blockade protocol was designed with the purpose of inhibiting the receptor, only during the specific postnatal period (P2-P14) when FLX induced depressive and anxiety-like phenotypes in the adult. This approach was shown to rescue the dysfunctional emotional behaviors in PNFLX-treated mice (41). Our findings in acute slices are consistent with our in vivo results, even though, due to the technical limitations of blind extracellular recordings in vivo, we cannot directly attribute the decreased firing rate, observed in intact animals, to the SERT+ subpopulation. Further experiments requiring opto-tagging of SERT+ PNs in vivo will be required to confirm that adult prefrontal hypoactivity resulting from PNFLX treatment is a specific attribute of this specific neuronal subtype.

Importantly, some studies in humans have shown that in utero exposure of SSRIs yields an increased risk to develop anxiety and depressive behaviors (69,70) and other cardiovascular anomalies (71). In this context, our findings could have translational importance when considering prescription of SSRIs during pregnancy and infancy.

Altogether, this study reports a marked effect on adult cortical network dynamics in mice, whose 5-HT levels were altered during an early-life sensitive period and resulting in depressive-like behavior. We identify a putative specific neuronal subtype, which seems to be particularly vulnerable to early-life insults, and we identify the 5-HT7R as a major molecular player. It will be interesting to identify how and when these alterations emerge during the development of cortical networks.

## Experimental Procedures

Experimental procedures followed national and European (2010/63/EU) guidelines and have been approved by the author’s institutional review boards and national authorities. All efforts were made to minimize suffering and reduce the number of animals.

### Animals

Mice were group-housed (3-5 per cage), maintained on a C57BL/6 background, and locally bred under standard laboratory conditions (22 ± 1°C, 60% relative humidity, 12-12 hrs. light–dark cycle, food and water *ad libitum*). In all experiments, both genders were equally represented. All neurophysiological recordings were performed on adult (>P90) mice. In order to record SERT+ pyramidal neurons we crossed B6.129(Cg)-*Slc6a4^tm1(cre)Xz^*/J (Sert-Cre) mice with B6.Cg-Gt(ROSA)26Sor<tm14(CAG-tdTomato)Hze>/J mice (td-Tomato) (The Jackson Laboratory, Bar Harbor). For in vivo experiments we used C57BL/6J mice (Janvier, France) and B6; 129S6-Htr7<tm1.2 Mrl> (5HT7 KO) mice (72).

### PNFLX protocol and drug administration

FLX (FLX, Tocris, UK; 10 mg/Kg) was diluted in a 2% sucrose solution and administered orally. Pups underwent a feeding protocol from P2 to P14, during which they were briefly separated from the home cage, weighed, and carefully fed with a micropipette either with the FLX/sucrose solution (FLX group) or with sucrose only solution (CTR group). SB269970 (here referred to as SB; 20 mg/kg; Abcam, #ab120508) or saline (0.9% NaCl) solution was injected intraperitoneally twice a day (every 12 hours) in FLX- and sucrose-fed pups (41). The entire procedure lasted <10 min per litter and had no visible consequences on maternal care to pups.

### In vivo electrophysiology

P90 – P105 mice, C57BL/6 or 5HT-7 KO of both sexes were anesthetized via intraperitoneal (i.p.) injection with 16% urethane (1.6 g/kg in 0.9% NaCl physiological solution) and placed on a stereotaxic apparatus. The body temperature was constantly monitored and kept at 37^◦^ C with a heating blanket. Eye ointment was applied to prevent dehydration. To ensure a deep and constant level of anesthesia, vibrissae movement, eyelid reflex, response to tail and toe pinching were visually controlled before and during the surgery. Subcutaneous injections of atropine (0.07 mg/kg) and glycopirrolate (0.01 mg/kg, Robinul-V, Vetoquinol) were used to maintain clear airways and prevent gasping. A subcutaneous lidocaine (Vetoquinol) injection was performed over the cranial area of interest then a longitudinal incision was done to expose the skull. A stainless-steel head post was sealed on to the mouse skull using dental acrylic cement. A small cranial window (<1 mm) was opened at coordinates (+ 2.5 mm to Bregma, 0.5 mm lateral to sagittal sinus), while keeping the surface of the brain moist with the normal HEPES-buffered artificial cerebrospinal fluid (aCSF).

For awake head-restrained recordings, mice were anesthetized with ketamine/xylazine mix (Ketaset, 100 mg/mL/Rompun 20 mg/mL; 10 mL/g body weight) to implant the stainless-steel head post as described above. Mice were kept at controlled body temperature until awakening. After 3 days of recovery, mice were habituated to the recording apparatus for at least 4 days with multiple sessions. The day of the experiment, a small (< 1mm) craniotomy was without removing the dura. The exposed cortical surface was superfused with warm HEPES-buffered extracellular solution (in mM: 125 NaCl, 5 KCl, 10 glucose, 10 HEPES, 1.8 CaCl2 and 1 MgSO_4_; pH 7.2) to maintain ionic balance and prevent desiccation. Electrophysiological juxtasomal recordings were obtained with borosilicate glass pipettes WPI (1.5 mm outer diameter, 5–9 MΩ resistance) pulled on a Narishige P100 Vertica Puller and filled with extracellular solution. To prevent tip occlusion, pipettes were initially lowered through the pia applying a high positive pressure (250 mbar) until the depth of interest was reached. At this point, a lower positive pressure (25 mbar) was preferred. Pipettes were advanced in 2-µm steps, and resistance was monitored in the conventional voltage clamp configuration. When the resistance slightly increased and a contact between the pipette and a putative cell was achieved, the positive pressure was released, and a dim negative pressure was applied to facilitate the formation of the juxtasomal recording configuration (resistance 40–400 MΩ). Recordings were made in current-clamp mode, and no holding current was applied. A typical recording duration lasted 10 min.

Data were recorded using a MultiClamp 700B amplifier, digitized with a Digidata 1440 and acquired using pClamp 10.7 software (Molecular Devices, USA). Raw data were sampled at 50 KHz and filtered at 10 KHz. Data were analyzed using custom-made scripts in Matlab (Mathworks, USA) and Python. Spikes were detected using a threshold algorithm for their time stamps. Juxtasomal recordings were high-passed filtered at frequencies larger than 300 Hz. The LFP signal was downsampled to 1 KHz and band-pass filtered between 0.1 and 200 Hz.

The temporal structure of spiking activity recorded from single mPFC neurons was analyzed to detect bursting activity using the Robust Gaussian surprise algorithm (73). The method consists of finding the subset of consecutive action potentials that have an associated probability of being observed that falls beyond a threshold value according to a normalized distribution. Briefly, inter-Spike Intervals (ISI) were computed for the spike train recorded from each neuron during 600 s of the total acquisition. The distribution of log(ISI) was computed and a Central Set of ISIs was defined as the set of log(ISI)s that fell within 1.64\**MAD* from the *E-center* where *MAD* is the median absolute deviation of log(ISI)s and *E-center* is the midpoint of log(ISI)s falling between the 5% and 95% quantiles. Normalized log (ISI) was calculated by subtracting the median of the Central Set. This normalization process was performed on the entire spike train using a sliding window of half-width 0.2*N/2, where N is the number of spikes in the spike train. The burst-threshold is set as 0.5 percentile of the central distribution. Pairs of action potentials were classified as burst seeds when normalized log (ISI) < burst-threshold and the number of action potentials belonging to the *k* burst (NB*_k_*) was set to 2 when the normalized ISI < burst-threshold, thus setting the burst seed (with 1 < k < total number of detected bursts). Each burst seed was then iteratively extended by including the subsequent normalized ISI of the spike train in the burst. The probability of observing NB*_k_* = 2 + *i* in the burst (P_NB*k*_) was computed for each iteration *i*. If the extension led to an increase of P_NB*k*_, the iteration was halted.Overlapping burst strings were cleaned by keeping the strings with the lowest associated *p*-value, thus making them mutually exclusive. Burst frequency was computed as the number of burst strings divided by the duration of the recorded spike trains.

Core code was taken from the Python implementation of the Robust Gaussian surprise algorithm made by Frontera et al. (74).

### In Vitro Slice Preparation and Electrophysiology

Coronal slices of the prefrontal cortex were obtained from adult (P90-105) SertCre-TdTom mice. Animals were deeply anesthetized with 16% urethane (SIGMA U2500) in 0.9% NaCl physiological solution. To ensure a deep and constant level of anesthesia, vibrissae movement, eyelid reflex, response to tail, and toe pinching were controlled before proceeding. After a lidocaine cream application (Anesderm 5%) at the incision site, animals were transcardially perfused using a cold (4°C) choline-based solution containing in mM: 126 choline chloride, 2.5 KCl, 1.25 NaH2PO4, 26 NaHCO3, 7 MgSO4, 0.5 CaCl2, 15 glucose, 3 pyruvic acid, 6 Myo-inositol, and 5 ascorbic acid, 2 thiourea (pH 7.3, 95% O2/5% CO2) and decapitated. The brain was quickly removed and immersed in cold choline-based solution (same used for the perfusion). Coronal slices (300 μm thick) were obtained from a block of brain that included the frontal lobes and were prepared with a vibratome (VT1200S, Leica Microsystems, GmbH, Wetzlar, Germany). Slices were incubated in oxygenated artificial cerebrospinal fluid (aSCF) containing the following in mM: 126 NaCl, 2.5 KCl, 2 CaCl2, 1 MgSO4, 1.25 NaH2PO4, 26 NaHCO3, and 16 glucose (pH 7.4), initially at 34°C for 15 min, and subsequently at room temperature for additional 30 minutes until transferred to the recording chamber.

Recordings were obtained at 32°C in the pre- and infralimbic areas of the PFC. Layer 5-6 SERT+ pyramidal neurons (labeled by TdTomato expression) were identified using LED illumination (green λ = 530 nm, OptoLED system, Cairn Research) coupled to the epifluorescent optical pathway of the microscope. SERT-negative pyramidal neurons were visually identified as tdTomato-negative neurons using infrared video microscopy and characterized by large somas with a pyramidal shape and the emergence of a thick apical dendrite oriented towards the pia.

Signals were amplified using a Multiclamp 700B patch-clamp amplifier (Molecular Devices, USA), sampled at 20 or 50 kHz and filtered at 4 kHz for voltage-clamp and 10 KHz in current-clamp mode. To record spiking activity, neurons were held in current-clamp mode, using a whole-cell pipette solution containing (in mM): 135 K-gluconate, 13 KCl, 10 HEPES, 1 EGTA, 2 MgCl2, 4 Mg-ATP, 0.3 Na-GTP,pH adjusted to 7.2 with KOH; 280-295 mOsm. Pipette resistance was 2-5 MΩ. Excitatory and inhibitory synaptic transmission was not blocked. Neurons were stimulated with 800s-long DC current injections starting at -100 pA and reaching 375 pA with current steps of 25 pA. Current injections were applied from resting membrane potential. The first negative current injection was used to measure sag and membrane resistance. Action potential firing and single-spike waveform were analyzed using a custom-written script in Matlab (Mathworks, USA).

For spontaneous glutamatergic excitatory postsynaptic currents (sEPSCs), the pipette solution was the same as that used for firing dynamics. SERT+ and SERT- PNs were held in voltage-clamp at -80 mV. sEPSCS were recorded over a period of 5-7 minutes immediately following whole-cell establishment. In some experiments, the ionotropic glutamate receptor antagonist 6, 7-dinitroquinoxaline-2,3-dione (DNQX, 10 µM) was applied and no remaining activity was recorded, confirming the glutamatergic nature of recorded sPSCs. For spontaneous inhibitory GABAergic postsynaptic currents (sIPSCs), the whole-cell pipette solution contained (in mM): 74 K-gluconate, 74 KCl, 3.2; 10 HEPES, 1 EGTA, 2 MgCl2, 4Mg-ATP, 0.3 Na-GTP, pH adjusted to 7.2 with KOH; 280-300 mOsm. This high-chloride intracellular solution yields a calculated equilibrium potential for Cl-mediated currents of ∼-15 mV when held at - 80 mV. In these experiments, glutamatergic transmission was blocked by the continuous presence of DNQX (10 µM) in the extracellular solution. In some experiments, to confirm the GABAergic nature of sIPSCs, gabazine (10 µM) was added to the aCSF. Recordings were discarded from analysis if access resistance exceeded 25 MΩ.

sEPSCs and sIPSCs were detected using custom-written software (WDetecta, courtesy J. R. Huguenard, Stanford University) (75,76). Briefly, individual events were detected with a threshold-triggered process from a differentiated copy of the real trace. For each cell, the detection criteria (threshold and duration of trigger for detection) were adjusted to ignore slow membrane fluctuations, artificial artifacts and electric noise while allowing maximal discrimination of IPSCs and EPSCs. Detection frames were regularly inspected visually to ensure that the detector was working properly. Population data were analyzed using Origin (OriginLab, USA), MATLAB (MathWorks), Python and GraphPad Prism software.

### Biocytin filling and neuronal reconstruction

To characterize the morphology of Layer 5/6 SERT+ and SERT- PNs, cells we performed whole-cell recordings of tdTomato-expressing and non-expressing neurons using an internal solution containing 5 mg/ml biocytin (Sigma). After at least 30 minutes recording, the patch pipette was gently removed until obtaining an outside-out patch. Slices were then fixed with 4% paraformaldehyde in phosphate buffer saline (PBS, Sigma) for 24 hr. Following fixation, slices were permeabilized in PBS-Triton 0.4%, then incubated in BSA 4%-PBS-Triton to block unspecified sites, and finally: incubate with Streptavidin (VectorLab Streptavidin DyLight® Alexa 488 1:1000). The following day, slices were subject to optical tissue clearing as in Meng-Tsen Ke and Takeshi Imai (77). Sections were then imaged in an Upright Spinning-disk confocal (3I/ZEISS) with a 25X objective by using SlidebookX64 acquisition software. Imaged neurons were then reconstructed and using Neurolucida 360 (MBF Bioscience) aided by the mouse atlas from the Allen Institute. Reconstructed neurons were analyzed in Neurolucida Explorer (MBF Bioscience). To determine changes in dendritic morphology, we performed Scholl analysis wherein number of intersections and/or total dendritic length intersection were calculated. Apical and basal dendrites were analyzed separately. A restricted maximum likelihood (REML) was performed to assess statistical differences between reconstructed cells of different types and in different treatments.

### Statistics

Values are expressed as mean ± SEM. To evaluate normality, a Shapiro normality test was run on each experimental dataset. When comparing 2 populations of data, Student’s *t*-test was used to calculate statistical significance in case of Gaussian distribution. In case of non-normal distributed datasets, the nonparametric Mann–Whitney test was used. When multiple populations of data were compared, one-way ANOVA with Bonferroni post hoc tests were used in the case of normally distributed datasets. Unless otherwise noted, for non-normally distributed multiple populations, non-parametric Kruskal– Wallis test was used followed by Dunn’s comparisons. All tests were two-sided.

## Acknowledgements

We thank the members of the Bacci laboratory for support, suggestions, and critical discussion. This work was supported by BBT-MOCONET1; BBT-MOCONET2; Agence Nationale de la Recherche ANR- 16-CE16-0007-02; ANR-17-CE16-0026-01; ANR-18-CE16-0011-01; ANR-20-CE16-0011-01; ANR-22- CE16-0021-02), Fondation Recherche Médicale (Equipe FRM DEQ20150331684 and EQU201903007860); Fondation Lejeune (#1790); ERA-Net NEURON Grant 221007_ANR ERA NET NEURON DevInDS. We thank the ICM technical facilities PHENO-ICMICE, iGENSEQ and ICM.Quant. PHENO-ICMICE was supported by two “Investissements d’avenir” (ANR-10-IAIHU-06 and ANR-11- INBS-0011-NeurATRIS). JZSM was funded by a postdoctoral research fellowship from Jerome Lejeune Foundation. NS is funded by the MSCA ITN network ‘Serotonin and Beyond’ GA-no. 953327.

## Author contributions

AMDS, PG and AB designed the research; AMDS and NS collected data. AMDS, JZDSM, NS and AB analyzed data; PG provided mouse lines; AMDS, JO, AA, JZDSM and NS performed the PNFLX feeding protocol; JL provided crucial feedback on electrophysiological experiments; AMDS and AB wrote the manuscript. All authors edited the manuscript.

## Supplementary Figures

**Suppl.FIG 1.**
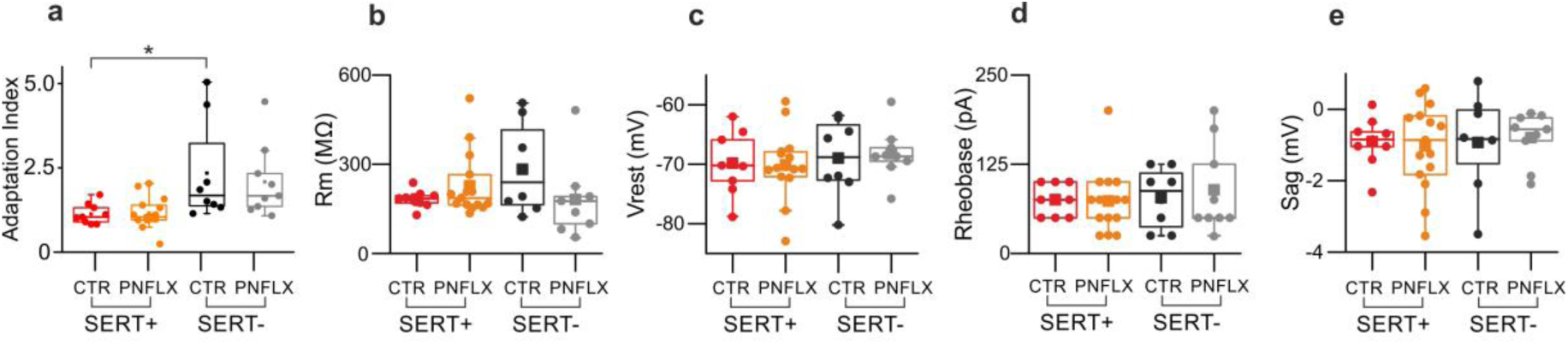
Adaptation Index and passive properties of SERT+ and SERT- PFC Pyramidal neurons in CTR and PFLX mice. **a-e:** Population data showing adaptation Index, membrane resistance, resting membrane potential, rheobase and Sag in computed for SERT+ and SERT+ cells in CTR (red and black) and PNFLX (orange and grey) mice.

**Suppl. FIG 2.**
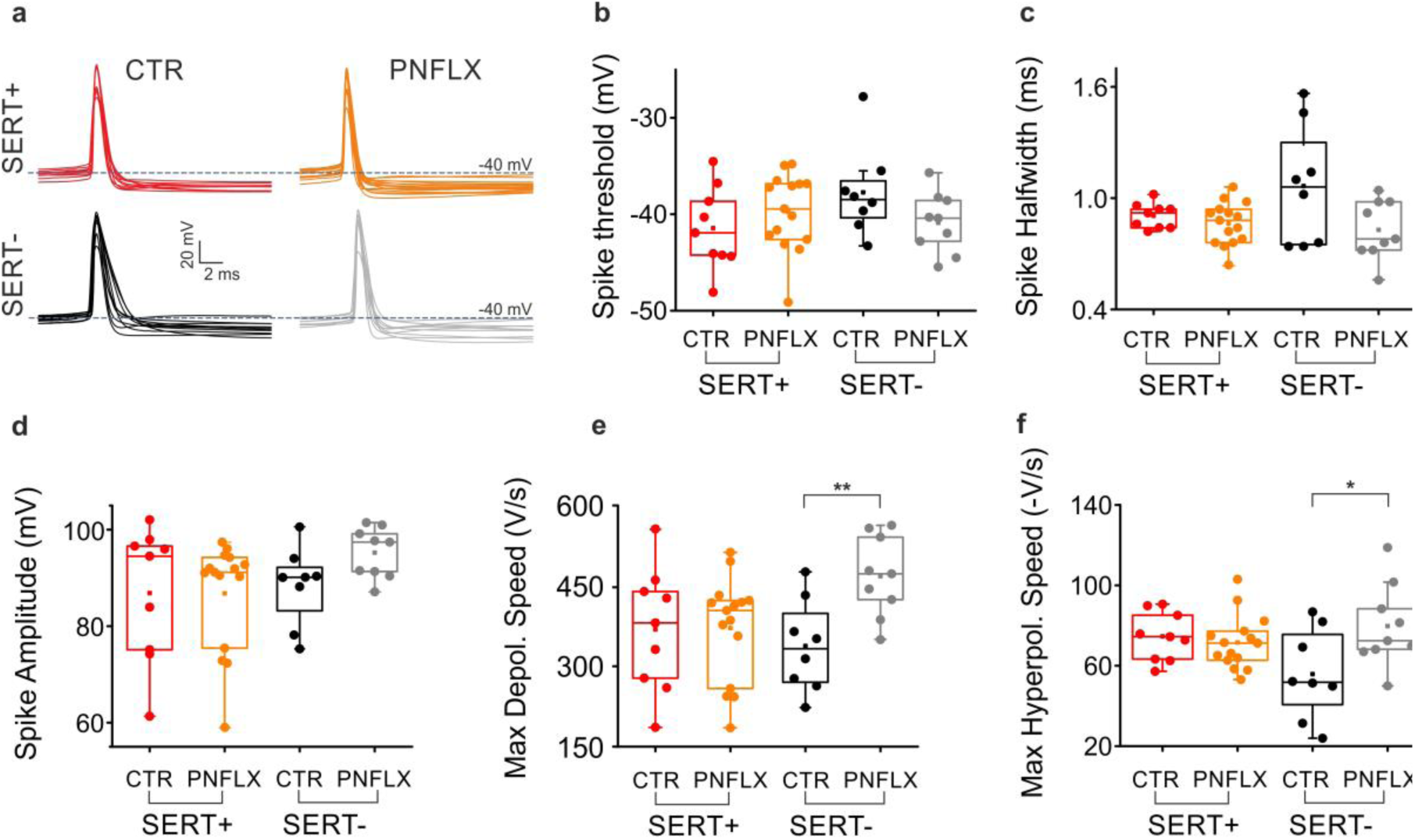
Action Potential properties in SERT+ and SERT- PNs. **a:** Representative single action potentials traces from all SERT+ and SERT- PNs in CTR and PNFLX treated mice. **b-f:** Population data relative to spike threshold, half-width, amplitude, max depolarization speed and max hyperpolarization speed in SERT+ and SERT- PNs, in CTR and PNFLX treated mice.

